# Piezo1 ion channels inherently function as independent mechanotransducers

**DOI:** 10.1101/2021.06.03.446975

**Authors:** Amanda H. Lewis, Jörg Grandl

## Abstract

Piezo1 is a mechanically activated ion channel involved in sensing forces in various cell types and tissues. Cryo-electron microscopy has revealed that the Piezo1 structure is bowl-shaped and capable of inducing membrane curvature via its extended footprint, which indirectly suggest that Piezo1 ion channels may bias each other’s spatial distribution and interact functionally. Here, we use cell-attached patch-clamp electrophysiology and pressure-clamp stimulation to functionally examine large numbers of membrane patches from cells expressing Piezo1 endogenously at low levels and cells overexpressing Piezo1 at high levels. Our data, together with stochastic simulations of Piezo1 spatial distributions, show that both at endogenous densities (1-2 channels/μm^2^), and at non-physiological densities (10-20 channels/μm^2^) predicted to cause substantial footprint overlap, Piezo1 density has no substantial effect on its pressure sensitivity or open probability in the nominal absence of membrane tension. The results suggest that Piezo channels, at densities likely to be physiologically relevant, inherently behave as independent mechanotransducers. We propose this property is essential for cells to transduce forces homogeneously across the entire cell membrane.

## Introduction

Cells are constantly exposed to mechanical forces and have evolved diverse molecular mechanisms that enable force detection. The rapid sensing of mechanical forces that occurs in milliseconds is achieved by force-gated ion channels that convert mechanical energy into electrochemical signals (Kefauver, Ward, and Patapoutian 2020). For example, Piezo1 is a mechanically gated cation channel that directly senses membrane tension (*T*) with high sensitivity (*T*_50_ ~ 1.4 mN/m) (Coste et al. 2010, Lewis and Grandl 2015, Cox et al. 2016, Syeda et al. 2016). Therefore, theoretically any mechanical disturbance of the cell membrane may lead to changes in Piezo1 channel open probability (P_o_). However, the propagation of membrane tension away from its source has been shown to be relatively slow (D ~0.024 μm^2^/s), poising Piezo channels with the ability to efficiently and rapidly transduce only mechanical forces that are nearby (Shi et al. 2018).

More generally, it can be said that the ability of a cell to transduce forces homogeneously across the cell membrane depends critically on *i)* the spatial distribution of force-gated ion channels, and *ii)* on their functional independence. Any substantial deviation from a uniform distribution of force-gated ion channels will result in domains that fall short to detect forces (where there are no ion channels) as well as domains that transduce forces disproportionately (where many ion channels are nearby). Functional interactions and/or cooperativity between force-gated ion channels may further skew transduction efficiency or kinetics in a manner that depends on channel density, effectively creating a patchwork of blind spots and sensitized areas.

Experimental evidence on the spatial distribution of Piezo1 ion channels is just beginning to emerge: TIRF imaging of Piezo1 channels revealed their free diffusion over the surface of live neural stem/progenitor cells, indirectly suggesting channels are not tethered to other static structures and that random membrane distribution may be possible. However, in the same cells, Piezo1-related activity was spatially enriched, which could arise from non-uniform distribution of Piezo1 channels, variable membrane tension, or both (Ellefsen et al. 2019). STORM imaging of HEK293 cells stably expressing GFP-tagged Piezo1 also suggests a non-homogenous spatial distribution, but at channel densities that are likely not physiological (Ridone et al. 2020). Finally, intensity distributions of GFP-tagged Piezo1 have been used to propose the existence of clusters with varying numbers of channels (Jiang et al. 2021).

In contrast, a compelling theoretical framework predicting spatial and functional interactions between nearby Piezo channels has been constructed based on cryo-em structures of Piezo1 and Piezo2, which revealed their large and extremely curved dome-shaped structure (Guo and MacKinnon 2017, Saotome et al. 2018, Haselwandter and MacKinnon 2018). Theoretical calculations based on these structures suggest that Piezo1 can both sense and curve the proximal membrane with a characteristic decay length extending ~14 nm beyond the channel. The energetic cost of this membrane curvature implies that Piezo1 channels that are within ~3 decay lengths (a “footprint” of ~50 nm; see **Methods**) may influence each other via multiple mechanisms: opposing curvature may cause nearby Piezo1 channels to repel each other, as well as influence each other’s gating properties (Haselwandter and MacKinnon 2018). Indeed, imaging of vesicles containing reconstituted Piezo1 have demonstrated the ability of Piezo1 to curve the membrane locally and suggested that flattening of the channel dome may directly couple to channel opening, but the precise effects of membrane energetics on gating have yet to be explored experimentally (Lin et al. 2019, Jiang et al. 2021). Here, we set out to precisely quantify the spatial distribution and potential functional interactions of Piezo1 ion channels across multiple orders of channel densities. Specifically, we use electrophysiology and large numbers of independent measurements to simultaneously quantify channel number and function, including single-channel current, open probability, and pressure-sensitivity, with a high level of precision. Collectively, our electrophysiological data, together with stochastic simulations show that Piezo1 ion channels, at densities likely to be physiologically relevant, inherently function as independent mechanotransducers. We propose this property enables cells to transduce forces with high spatial homogeneity across the entire cell membrane.

## Results

### Piezo1 function at low channel densities

To simultaneously measure and thus directly correlate Piezo1 channel density with its gating properties, we performed cell-attached patch-clamp electrophysiology on Neuro2A cells, which natively express Piezo1 (Coste et al. 2010). We chose Neuro2A cells over human umbilical vein endothelial cells (HUVECs; **Figure S1A-C**) and other cell lines because they have among the highest endogenous levels of Piezo1 expression, thus offering the highest likelihood for observing any potential functional effects of local channel density on function. We then designed a novel stimulation protocol that we optimized to assess Piezo1 single-channel conductance (i), pressure sensitivity (P_50_), and number of channels (n) in each patch with high accuracy. Since Piezo1 inactivation precludes a precise measurement of maximal current (I_max_) at negative potentials, we recorded currents at +60 mV, where inactivation is minimal (Coste et al. 2010, Wu et al. 2017). However, traditional step protocols are well-known to induce irreversible changes in patch geometry owing to membrane creep, particularly at positive voltages (Suchyna, Markin, and Sachs 2009, Lewis and Grandl 2015). Indeed, a direct step to a saturating pressure (−60 mmHg) caused substantial changes in current due to leak and/or capacitance even in patches that were later classified to have zero channels (see below; mean±S.D.: 2.6±1.3 pA; n = 33 patches; **Figure S2A-B**). Consistent with this idea, the same pressure step caused a similarly small, but measurable current in patches from Neuro2A-Piezo1ko cells, which have been CRISPR-engineered to lack Piezo1 channels (mean±S.D.: 3.8 ±1.5 pA, n = 15 patches; **Figure S2C**). Consequently, a step protocol consistently overestimates current and thus channel numbers (**Figure S2B**).

To overcome this limitation, we instead designed a protocol in which the pressure was decreased incrementally in 1 mmHg steps from 0 mmHg to −60 mmHg in 50 ms intervals, thus approximating a ramp (**Figure 1A-D**). The total duration of this ramp is short enough to minimize membrane creep (3 s), but each step is long enough (50 ms) to allow pressure equilibration, which occurs within ~7 ms (Lewis et al. 2017). Similar to the step protocol, the ramp protocol elicited small background changes in current amplitude in all patches. However, instead of the square-like current increases evoked by the step protocol, here the background changed more steadily (**Figure 1A-D, middle**), allowing us to perform linear background subtractions for each individual patch and ultimately achieve a more reliable detection of channel gating events, which were easily discerned as nearly instantaneous changes in current amplitude (**Figure 1A-D, bottom**). We are confident that the slow changes in current are unrelated to Piezo gating, as we never observed channel openings during the ramp phase of the protocol in 15 patches from Neuro2A-Piezo1ko cells (data not shown). Ultimately, we decided to use the ramp stimulus protocol followed by a brief (200 ms) saturating pressure step, the latter of which allowed us to confirm post-hoc that inactivation of Piezo1 at these voltages is minimal (step pulse: mean current at 200 ms = 92.5±14.2% of peak; n = 281 patches; **Figure S2A**).

**Figure 1:**
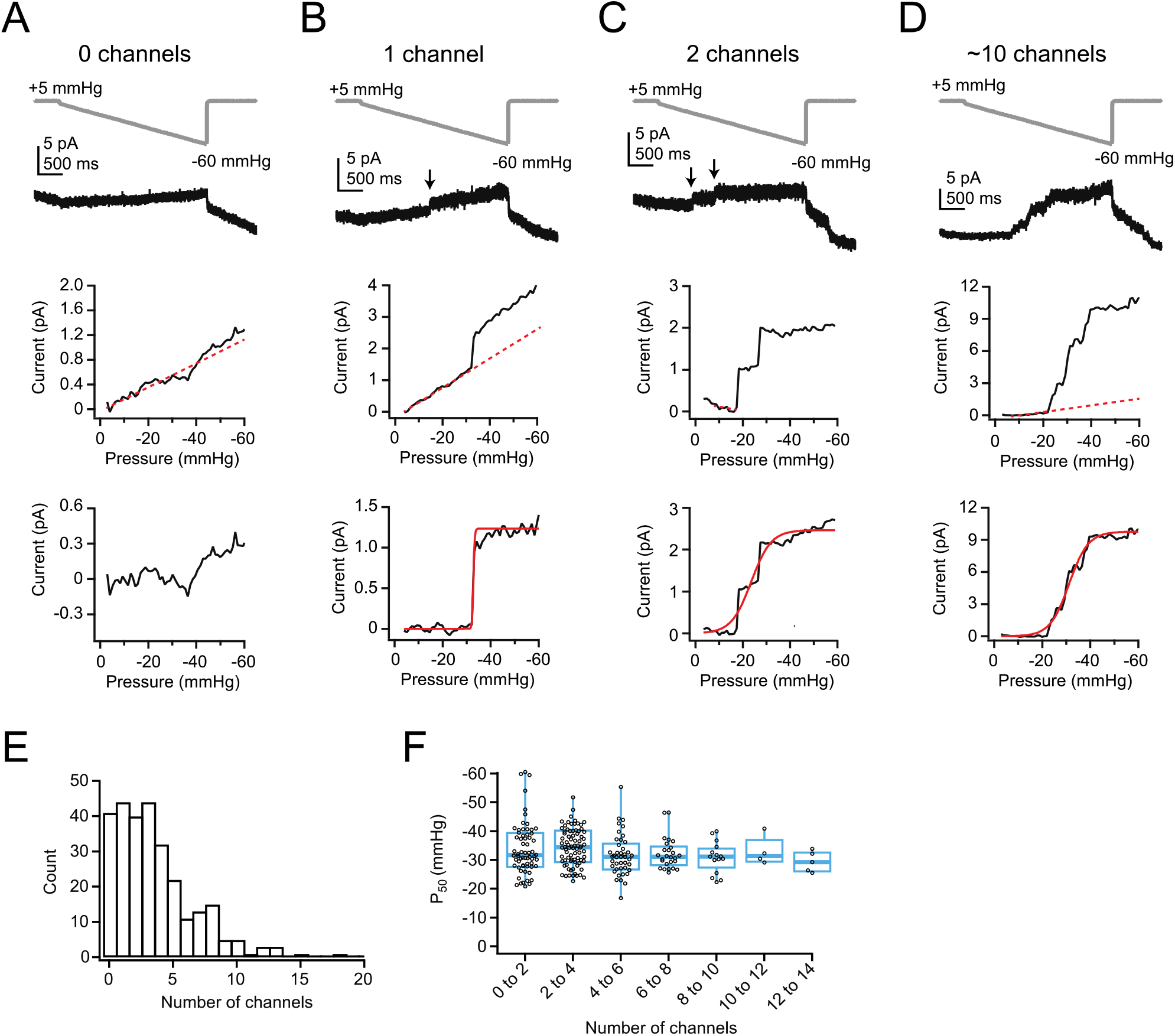
Low levels of Piezo1 expression in Neuro2A cells do not influence pressure sensitivity. **A**, *Top*, Pressure protocol (gray) and current (black) from a cell-attached patch from a Neuro2A cells with 0 channels. Holding potential was +60 mV. *Middle,* Current-pressure relationship for the same patch. Mean current was calculated for each step, in 1 mmHg increments, and plotted as a function of pressure. Note the linear relationship between current and pressure during the step, despite no channel activity in this patch (inferred from the lack of discrete channel opening events). This is likely due to small changes in seal quality and/or capacitance. The linear portion of the current-pressure relationship was fitted with a line (red dashes). *Bottom*, Current as a function of pressure after subtraction of the current corresponding to the fitted line. **B**, As in A, for a patch with 1 channel. Arrow indicates discrete channel opening event. *Bottom,* current was fitted with a Boltzmann equation (see methods). From the fit (red line), I_max_, which corresponds to single-channel current, was 1.2 pA and P_50_ was −32.8 mmHg. **C**, As in A-B, for a patch with two channels. From the fit (red line), I_max_ was 2.5 pA and P_50_ was −23.6 mmHg. **D**, As in A-C, for a patch with many channels. From the fit (red line), I_max_ was 9.8 pA and P_50_ was −31.2 mmHg. **E**, Histogram of channel number in each patch. n = 281 patches. **F**, P_50_ values, measured from sigmoidal fits to currents in A-D, as a function of channel number. Bin width is 2 channels. n = 281 patches.

Next, we focused on titrating the sizes of our patch pipettes such that a substantial fraction of patches had zero channels (**Figure S2D**; see Methods for details). We limited our pipettes to 3-6.5 MΩ reasoning that this choice maximizes our ability to detect potential channel clusters and thus resolve the overall spatial distribution of channels. Also, to address the possibility that Piezo1 cellular distribution could be altered by the cell-substrate interface, which we reasoned may trap Piezo1 channels and deplete them from cell-attached patch-recordings on the cell roof, we performed day-matched experiments of Neuro2A cells that had either been seeded to adhere overnight or were acutely replated <1 hour before recording, therefore allowing limited time for channel redistribution (see Methods). Importantly, we found that the mean channel number and distributions in these two datasets were identical (mean± S.E.M: overnight = 2.4±0.4 channels, n = 28 patches; acute = 2.7±0.4 channels, n = 26 patches, p=0.6, Student’s t-test; **Figure S2E**), which argues against an effect of channel redistribution to the cell-substrate interface. Finally, in order to obtain good representations of the spatial distribution and function of Piezo1 channels, we executed this protocol on patches from a very large number of individual Neuro2A cells (n = 281).

After background subtraction, we could easily recognize in most patches (248/281) discrete, nearly instantaneous increases in current amplitude that are reflective of individual channel openings, as well as clearly identify a plateau phase of the current (**Figure 1B, middle**). The remaining patches (33/281) were classified as having zero channels (e.g., **Figure 1A**). We next used the mean single channel current from all patches with one channel (i = 0.98±0.04 pA, n = 35 patches; **Figure S2F**) to calculate the precise number of channels (n = I_max_/i) in all other patches. Ultimately, we obtained a continuous, bell-shaped distribution of Piezo1 channel numbers per patch (**Figure 1E**). The distribution average was 3.5±3.1 channels per patch, which given our small pipette sizes is consistent with previously reported levels of expression in Neuro2A cells (Coste et al. 2010). We also applied each protocol twice per patch and then performed pairwise comparison of current amplitudes (**Figure S2G-H**). This analysis revealed an extremely low variability (ratio of 2nd/1st recording = 1.1±0.4; n = 261), supporting that the assessment of channel numbers is highly accurate. We then used an estimate of the dome size of membrane patches inside patch-pipettes from imaging experiments we performed previously in our lab ((Lewis and Grandl 2015); **Figure S3**; see **Methods**) to calculate the average density Piezo1 channels to be 1-2 channels/μm^2^.

Next, we used two different approaches to assess if density of Piezo1 channels affects their sensitivity to membrane tension: First, we calculated P_50_ values, measured from sigmoidal fits to the ramp phase of the current, and plotted them as a function of channel number (**Figure 1F**). Although P_50_ measurements varied widely in individual patches, especially for those with <5 channels, because of the stochastic nature of channel gating (e.g., **Figure 1C**), this variance is partially overcome by the large number of recordings we performed (n = 248). Importantly, while there was some variability between two protocols executed on the same patch, there was only a slight systematic shift in P_50_ values toward smaller pressures in the second protocol, likely due to creep and corresponding increase in patch radius (mean±S.D. first = −33.1±7.2 mmHg, second = −31.1±7.7 mmHg; ΔP_50_ = +2.0±8.2 mmHg; p<0.05, paired t-test; **Figure S2I**), providing further evidence that our protocol allowed for highly reproducible measurements. Altogether, the results show that, for patches containing 1-15 channels, average P_50_ values do not vary substantially with channel number (**Figure 1F**).

Still, to measure P_50_ values in patches with few channels more precisely, we also performed a different analysis: Specifically, we idealized recordings with ≦5 channels by identifying discrete openings (**Figure 2A-E; see Methods**) and then averaged all traces with equal channel numbers in order to obtain their pressure-response curves (**Figure 2B**). Visual inspection as well as fits with sigmoidal functions show that P_50_ and slope (k) values are virtually identical for patches containing 1-5 channels (1 channel: P_50_ = −23.2 mmHg, k = −3.3 mmHg; 5 channels: P_50_ = −23.7 mmHg, k = −4.2 mmHg). We therefore conclude from both analyses that at densities of up to 2 channels/μm^2^, Piezo1 channels do not influence each other’s mechanosensitivity.

**Figure 2:**
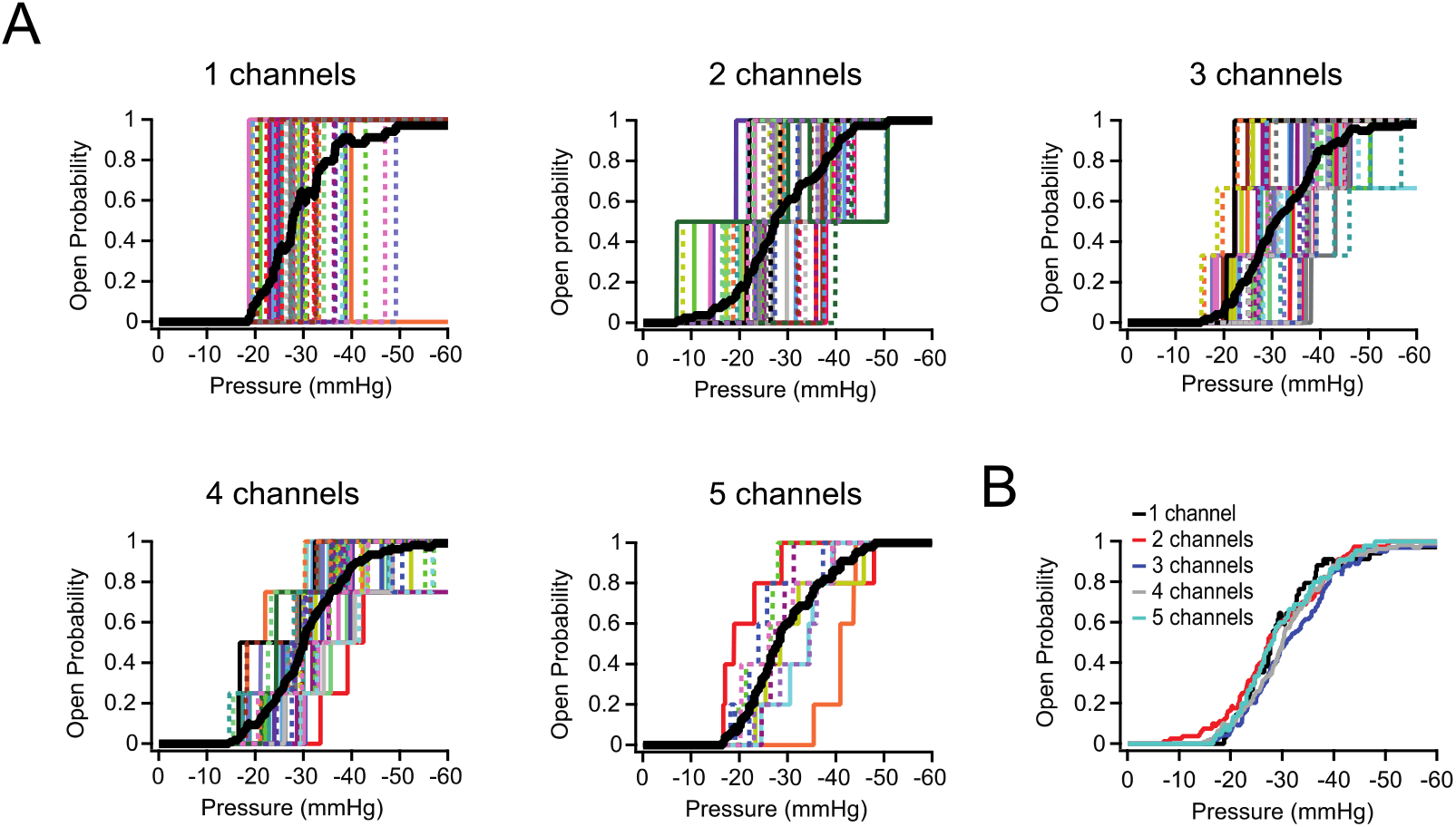
P_50_ and slope values are identical for Neuro2A patches with 1-5 Piezo1 channels. **A**, Schematized current-pressure relationship for all patches with precisely 1-5 channels. Each patch is a different color/weight; thick black line is the mean open probability for all patches. 1 channel: P_50_ = −23.2 mmHg, k = −3.3 mmHg, n = 35 patches. 2 channels: P_50_ = −23.6 mmHg, k = −5.1 mmHg, n = 40 patches. 3 channels: P_50_ = −25.9 mmHg, k = −4.5 print mmHg, n = 33 patches. 4 channels: P_50_ = −24.9 mmHg, k = −4.0 mmHg, n = 27 patches. 5 channels: P_50_ = −23.7 mmHg, k = −4.2 mmHg, n = 9 patches. **B**, Overlaid mean open probability as a function of pressure for patches with precisely 1-5 channels.

### Spatial distribution of Piezo1

Multiple mechanisms may explain our above result that Piezo1 pressure sensitivity does not vary with channel density. First, Piezo1 channels may be randomly distributed and therefore spatially too distant to functionally interact via their membrane footprints. Alternatively, Piezo1 channels may indeed tend to localize in close spatial proximity (i.e., groups of 2-3 channels), but this localization may have no effect on their pressure sensitivity. To begin distinguishing among these possibilities, we built a statistical model for spatial localization, based on a Thomas point process (see Methods for details). First, we simulated a distribution of randomly dispersed Piezo1 channels. Then we randomly sampled from this population, accounting for variability of pipette sizes and single-channel currents by using our experimental mean values and standard deviations of pipette size and single-channel conductance (**Figure S2D,F**). The resulting distribution fits the experimental data reasonably well, indicating that in Neuro2A cells, Piezo1 channels may be homogenously distributed across the cell surface, rather than being spatially localized in close proximity (**Figure 3A-D**).

**Figure 3:**
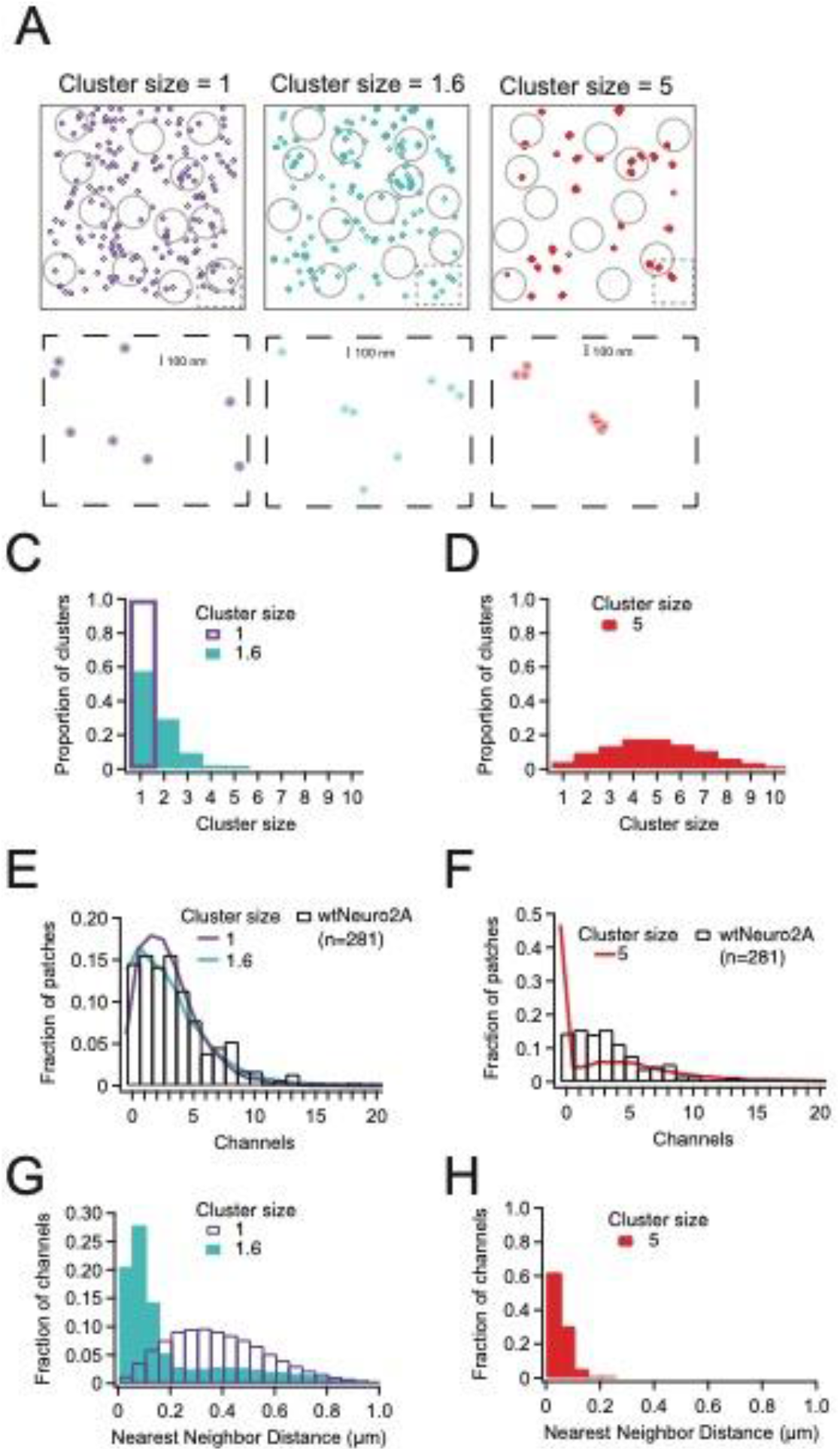
Piezo1 distribution in Neuro2a cells is best explained by little or no clustering of channels. **A**, Representative channel distributions generated using a Thomas point process with overall densities equivalent to that of wild-type Neuro2A cells (~1.75 channels/μm^2^); in each condition, daughter “channels” are assigned to a center “parent” disc, with a mean distance of 50 nm (~one Piezo footprint) from the center of the disc. Three separate degrees of clustering (“cluster size”) were introduced; one in which every cluster has precisely 1 channel, one with a mean of 1.6 channels per cluster, and one with a mean of 5 channels per cluster. Gray circles represent model patch domes used to sample the distributions, with a mean radius of 0.8 μm. Below, insets (dashed boxes) of each distribution; dark circles are the projected area of the Piezo footprint to scale; light circles are ~3x the membrane footprint for each channel (radius = 50 nm). **C-D**, Distributions of channels per cluster for mean cluster sizes of 1 and 1.6 (C) and 5 (D). **E-F**, Normalized histogram of channels per patch for wild-type Neuro2A cells for mean cluster sizes of 1 and 1.6 (E) and 5 (F). **G-H**, Histogram of nearest-neighbor distances for mean cluster sizes of 1 and 1.6 (G) and 5 (H).

In order to challenge this interpretation, we next simulated a random distribution of Piezo1 clusters in which individual Piezo channels are distributed across a 2D-Gaussian distribution with a standard deviation of 50 nm which is our estimate for the Piezo channel footprint, all while holding the overall channel density constant. Strikingly, a distribution in which on average 1.6 Piezo1 channels (drawn from a Poisson distribution centered at 1; see **Methods**) interact over 50 nm also describes the experimental data well. However, simulating larger channel clusters (n ~ 5) fails to reproduce the experimental distribution, specifically the probability of capturing patches without any channels (**Figure 3A-E**). Thus, our experimental data are most parsimoniously explained by either a random spatial distribution or a weak propensity for Piezo channels to spatially localize in groups of 2-3. Interestingly, the two distributions differ significantly in the predicted fraction of channels that are within one footprint of at least one other channel: in a truly random distribution, the mean nearest neighbor distance is ~390 nm, and only ~5% of channels are within 100 nm of another channel, whereas if channels tend to cluster in groups of 2-3, the mean nearest neighbor distance is reduced to 225 nm and almost 50% of channels are within 100 nm of another channel (**Figure 3F**). However, our functional data support the idea that if the latter spatial localization indeed occurs, it does not have a substantial effect on Piezo1 pressure sensitivity (**Figures 1 and 2**).

### Piezo1 function at high channel densities

Our above results do not answer if, in principle, Piezo1 channels have the ability to functionally interact. We therefore decided next to examine Piezo1 function using a heterologous overexpression system, in which channel densities are much higher. For these experiments, we chose Neuro2A-Piezo1ko cells, in order to retain the same cellular background, but avoid heterogeneity from endogenous and overexpressed Piezo1, especially given that there are known splice variants of Piezo1 (Geng et al. 2020). In pilot experiments, we found that transfection with Lipofectamine 2000 maximizes channel density 42-48 hours post-transfection (data not shown). Since overexpression yields I_max_ currents that are much larger, but also causes a higher variability in the degree of inactivation even at positive potentials (**Figure S4A-C**; current at 200 ms = 53.3±18.1% of peak, n = 138 patches), this time we chose a classic pressure-step protocol to best measure both peak current (I_max_) and pressure-sensitivity (P_50_ and k). While the step protocol still produces a 2-3 pA background in peak amplitude, in the overexpression system this error is very minor compared to the generally large currents, especially for patches with many channels. To reduce bias from resting membrane tension, each pressure step was preceded by a 5 second +5 mmHg prepulse (Lewis and Grandl 2015), and to minimize the effects of repetitive stimulation on membrane creep, we recorded at −80 mV and limited each step to 250 ms. We also used a pressure increment (ΔP) of −5 mmHg, so that we could better resolve small changes in pressure sensitivity (**Figure 4A-B**). As before, we restricted our pipettes to 3-6.5 MΩ, a range in which peak current amplitudes do not vary substantially (**Figure S6A**). We again repeated this measurement for many individual patches (n = 133 patches) in order to capture an accurate representation of channel densities and functional interactions.

**Figure 4:**
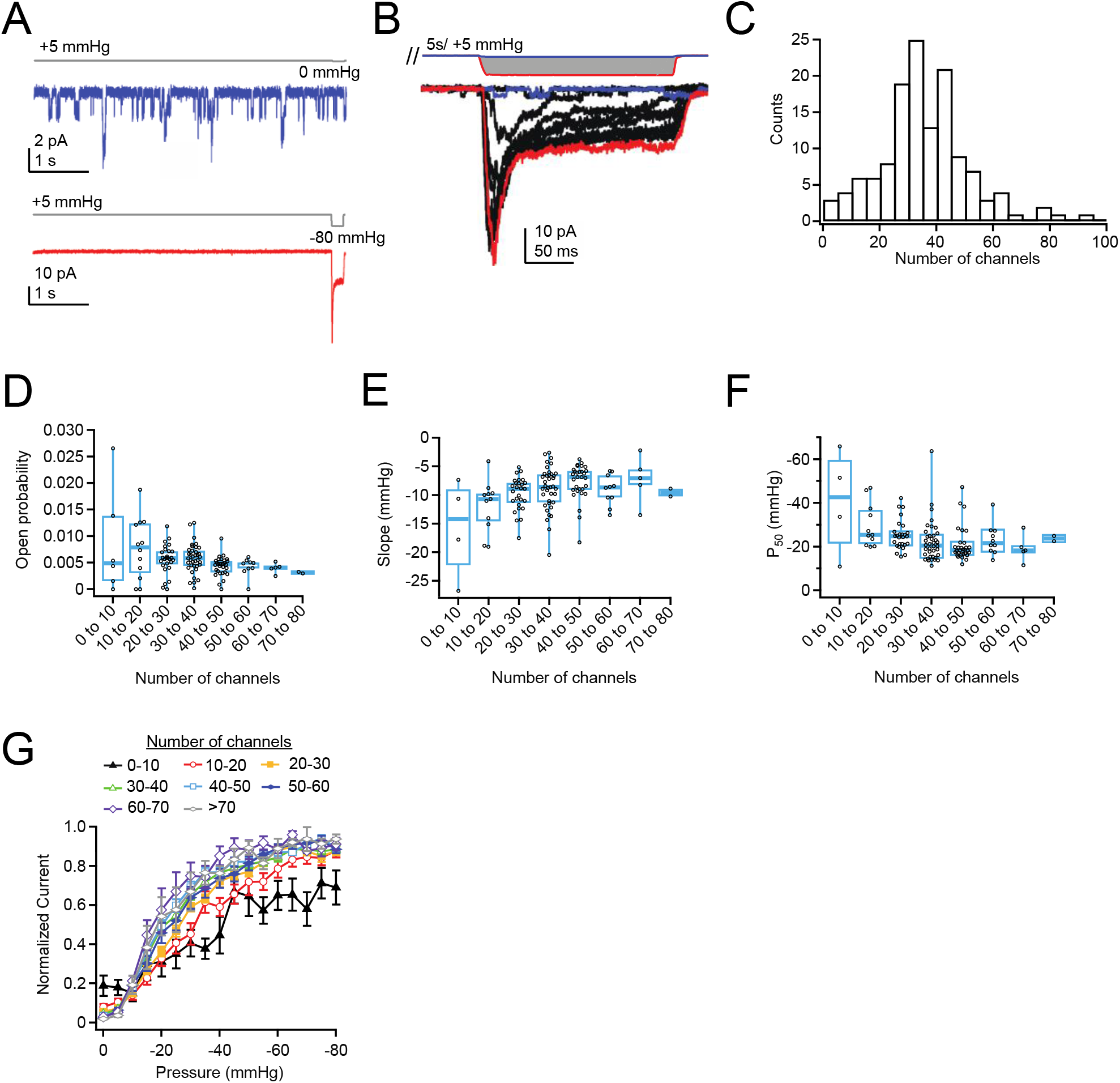
Density of overexpressed Piezo1 channels does not affect resting open probability or pressure sensitivity. **A**, Pressure protocol (gray) and current from a cell-attached patch from a Neuro2A-p1ko cell overexpressing mouse Piezo1. Pressure-steps were to 0 mmHg (blue) and −80 mmHg (red) and were preceded by a 5 s prepulse to +5 mmHg to relieve resting membrane tension. **B**, Full pressure-response protocol and currents for the cell in A. Pressure steps were from 0 to −80 mmHg in −5 mmHg increments. **C**, Histogram of channel counts, obtained by dividing the peak current in B by the single channel current measured from unitary events during the prepulse in A (see also **Figure S5**). n = 133 patches. **D**, Open probability during the +5 mmHg prepulse as a function of number of channels. n = 2-38 patches. **E**, Slope (k) values measured from sigmoidal fits to current-pressure relationships from the protocol in B as a function of the number of channels. **F**, P_50_ values measured from sigmoidal fits to current-pressure relationships from the protocol in B as a function of number of channels. **G**, Mean pressure-response curves, generated by first normalizing each patch to its maximal current, binning by number of channels, then averaging normalized currents at each pressure. n = 2-38 patches.

The low channel activity during the prepulse allowed us to measure the single-channel current of each individual patch (**Figure S5**) and therefore to precisely calculate the minimal number of channels (n = I_max_/i). This is particularly important in a cell-attached configuration, where even in a high K^+^ solution, there can be slight variance in the resting membrane potential and correspondingly variance in the single-channel current in each cell that introduces an additional source of error. Importantly, there was no correlation between number of channels and single-channel current, suggesting that unitary conductance of Piezo1 is not modulated by channel density (**Figure S5C**). The number of channels per patch was well-described by a Gaussian distribution with 35.3±15.9 channels per patch (**Figure 4C**), which we estimate corresponds to a Piezo1 density of 10-20 channels/μm^2^ (see Methods). The most extreme current amplitude we observed (−160 pA, or ~80 channels) corresponds to a maximal density of ~40 channels/μm^2^. At this maximal density, if Piezo channels were randomly distributed in the membrane, their mean next-neighbor distance would be ~80 nm, and 31% of the membrane surface would be covered by Piezo proteins and their corresponding membrane footprints (**Figure S7**). This calculation suggests that we have reached densities in which Piezos are likely in close enough proximity to influence each other’s function. Moreover, this calculation likely represents a slight underestimate of the true channel density, owing to inactivation at negative potentials. Two groups had previously predicted that nearby Piezo1 channels may influence each other’s gating, even in the absence of membrane tension (Haselwandter and MacKinnon 2018, Jiang et al. 2021). We therefore chose first to analyze the activity during the prepulse, which is designed to minimize resting membrane tension. We found that open probability was generally small (~1%), which was consistent with previous work in our lab (Wu, Goyal, and Grandl 2016). Most notably however, we found that Piezo1 open probability did not increase with channel density; rather, open probability slightly decreased, from 0.86±1.00% for patches with 1-10 channels (mean ± S.D.; n = 6 patches) to 0.40±0.09% for patches with 60-70 channels (n = 5 patches) (**Figure 4D**). There was no change in open probability with pipette resistance, indicating that variability in the ability of the prepulse to minimize resting tension with pipette size does not account for this trend in open probability with channel number (**Figure S6B**).

As an alternative test, we measured the effect of channel number on pressure sensitivity (P_50_ and slope (k)). The results show that for patches containing >20 channels, neither slope nor P_50_ values are affected by channel number (**Figure 4E**). The average P_50_ value for <20 channels appears slightly right-shifted, but is also relatively imprecise, because the stochastic nature of channel gating makes pressure-response curves noisy and hard to capture with sigmoid fits, and efficient averaging is precluded by the few patches with low channel numbers. Therefore, we also analyzed the data in a separate way: We normalized each patch to its maximal current, binned patches according to their number of channels, and then averaged pressure-response curves (**Figure 4F**). The results show at most a slight left-shift in pressure-response curves and increase in sensitivity (slope) with increasing channel number (for 20-30 channels, P_50_ = −24.1 mmHg, k = −9.9 mmHg, n = 27; for 60-70 channels, P_50_ = −17.1 mmHg, k = −6.7 mmHg, n = 10). Altogether, we conclude that at high densities, Piezo1 channels may be slightly more sensitive to pressure, but that there is no positive cooperativity in the nominal absence of membrane tension.

## Discussion

We sought to characterize the inherent ability of Piezo channels to spatially localize and modulate each other’s function. We chose cell-attached patch-clamp electrophysiology as a method for investigation, which has several inherent limitations that need to be considered. First, our spatial resolution is naturally limited by the size of the patch pipette. Consequently, our statistical approach of counting channels within membrane patches works best when the spatial distance between Piezo1 channels is comparable to the size of the membrane dome inside the patch pipette. The fact that ~15% of patches in wild-type Neuro2A cells contained no Piezo1 channels (**Figure 1E**) is therefore an extremely important quality benchmark for adequate spatial resolution. Our modeling does not allow us to rule out the possibility of moderate clustering (2-3 Piezo1 channels), but we find no evidence for clusters of larger sizes (~5 channels), which we would have been able to detect.

Next, our measurement of endogenous Piezo1 density in Neuro2A cells (1.75 channels/μm2) has several sources of error: First, while we have modest control over the pipette shape during its fabrication and a precise measure of its resistance, the exact surface of each membrane dome is not known. We returned to previous experiments, where we optically determined the geometry of membrane domes inside patch pipettes, to estimate patch dome area, but the drastically limited throughput of that technique precluded individually measuring each pipette (Lewis and Grandl 2015). In order to minimize this size variability, we therefore limited our analysis to measurements from a narrow range of pipette resistances and incorporated this size variability of patch pipettes into our statistical model of Piezo1 spatial distribution (**Figure S2D**). Certainly, when overexpressing Piezo1, the variability in Piezo1 membrane levels is high, in part due to variable expression level among cells. It is also subject to additional error due to the presence of inactivation, which prevents observation of P_o_ = 1. However, none of the above statistical errors, or others we did not consider and that skew channel number distributions, affect our ability to correlate Piezo1 channel number with its function.

The weaknesses of cell-attached patch-clamp electrophysiology are balanced by two clear strengths: the method has relatively high throughput so that large numbers of independent measurements yield good population averages, and it has unmatched precision: channel number and function, including single-channel current, open probability and pressure-sensitivity can be extracted over nearly two orders of magnitude (1-80 channels) with high accuracy. For example, repeated execution of pressure ramps on each patch revealed that channel number (n) is measured with only 24% variability, demonstrating that this assessment is extremely accurate.

Neither endogenous Piezo1 in Neuro2A cells or overexpression of Piezo1 in Neuro2A-P1ko cells are perfect model systems for investigating how Piezo1 channels function when they are directly contacting each other: experimental channel densities do not reach the theoretical limit of a tightly packed 2D hexagonal lattice. However, the maximal densities we obtained are certainly sufficient for Piezo1 membrane footprints to geometrically overlap: for example, at the highest density we measured (~80 channels/μm^2^), the centers of randomly spaced Piezo1 domes are on average 80 nm from their nearest neighbor, which results in a substantial number of channels having overlapping membrane footprints (Haselwandter and MacKinnon 2018). Additionally, in the overexpression experiments our step protocol may slightly underestimate channel numbers, as inactivation at negative potentials likely precludes piezo1 from reaching maximal open probability. In any case, two distinct analyses show that at densities of 1-2 channels/μm^2^ (**Figure 1-2**) and 10-20 channels/μm^2^ (**Figure 4**) Piezo1 pressure sensitivity is not modulated by channel numbers. A mechanistic explanation for this result at low densities is directly provided by our data and statistical model of spatial distributions, which suggest that in Neuro2A cells, Piezo1 channels are either randomly distributed or have at best a very low probability of aggregating. This is consistent with Haselwandter and MacKinnon’s prediction that the Piezo structure favors channels to repel each other. However, the result does not mean that Piezo1 channels in close proximity can never be found. A mild tendency for clustering predicts 20% of all Piezo1 channels to be within 50 nm of each other (cluster size = 1.6; **Figure 3F**), which is fully consistent with studies that have captured images of neighboring Piezo1 channels (Ridone et al. 2020), but 80% of all Piezo1 channels are outside this range. In other words, in wild-type Neuro2A cells, most Piezo1 channels are too far apart to ‘feel’ each other. As a consequence, they behave as functionally independent mechanotransducers. This result is important, because it shows that Piezo1 channels have no strong intrinsic tendency to spatially concentrate and functionally modulate each other’s mechanical sensitivity. Both of these properties are ideal for a cell in order to transduce mechanical forces homogeneously. Still, does the Piezo1 membrane footprint allow for the possibility of cooperativity among nearby channels? Our experiments at high channel densities (10-20 channels/μm^2^) reveal at most a modest sensitization of Piezo1 pressure sensitivity with increasing channel numbers. While this result supports the idea of an interaction via the membrane footprint other mechanisms may well be responsible and must be considered: for example, calcium permeating through Piezo1 may activate or sensitize nearby channels. Indeed, calcium-induced cooperativity is a well-known phenomenon in other ion channels, such as TRPA1 (Wang et al. 2008). Our experiments cannot directly rule out this explanation.

Surprisingly, our data suggest that in the nominal absence of membrane tension Piezo1 open probability is slightly reduced with channel number and overall, very low (< 1%) at all channel densities we probed. This is notable, because the opposite result was recently obtained by simulations of Piezo1 that are in extreme proximity to each other (1-3 nm dome-dome distance; (Jiang et al. 2021)). The reasons for this discrepancy may be technical limitations in the modeling, which was performed with a partial Piezo1 structure, or our own experiments, which cannot generate the extreme channel densities probed in the simulation.

Altogether, our conclusions may be applicable to other cell types that express Piezos at levels similar or lower to that of Neuro2A. However, we fully expect that specific cell types may concentrate Piezo channels into locations that are dedicated to detecting forces. For example, it is plausible that in sensory neurons Piezo2 channels may be highly concentrated in free nerve endings as compared to the cell soma. However, our work directly predicts that if such a clustering was indeed observed, it would require a dedicated mechanism that is not Piezo-intrinsic, such as a still elusive tethering molecule.

## Acknowledgements

This study was supported by NIH 5R01NS110552 (A.H.L. and J.G) and Duke University. We thank Gary Lewin for generously providing Neuro2A-P1ko cells, and all members of the Grandl lab for thoughtful comments on the study.

## Methods

### Cell Culture

Wild-type Neuro2A cells (ATCC #CCL-131) and Neuro2A-p1ko cells (a gift of Gary Lewin) were cultured at 37°C and 5% CO_2_ in Minimum Essential Medium (ThermoFisher Scientific) supplemented with 0.1 mM non-essential amino acids, 1 mM pyruvate, 10% FBS (Clontech), 50 U/mL penicillin, and 50 mg/mL streptomycin (Life Technologies). Human umbilical vein endothelial cells (HUVECs; Lonza CC-2519, pooled) were cultured in EGM-1. For experiments without transfection, cells were directly plated on poly-L-lysine and laminin-coated coverslips 16-24 hours before recording. For overexpression experiments, cells were transiently transfected with 4 μg mouse Piezo1-pIRES-EGFP in pcDNA3.1(+) or 4 μg YFP 40-48 hours before recording in 6-well plates using Lipofectamine 2000 (Thermo Fisher Scientific) according to the manufacturer’s protocol. Transfected cells were reseeded onto poly-L-lysine and laminin-coated coverslips 16-24 hours before recording. Only one patch was made from each cell; in some cases, two protocols were run on each patch (as noted). For acute plating experiments, cells were reseeded onto poly-L-lysine and laminin-coated coverslips, allowed to settle for 45 minutes, and then recorded for up to one additional hour.

### Electrophysiology

Electrophysiological recordings were performed at room temperature using an EPC10 amplifier and Patchmaster software (HEKA Elektronik, Lambrecht, Germany). Data were sampled at 5 kHz (wild-type Neuro2A) or 10 kHz (Neuro2a-p1ko + mPiezo1) and filtered at 2.9 kHz. The cell-attached bath solution used to zero the membrane potential was (in mM): 140 KCl, 10 HEPES, 1 MgCl_2_, 10 glucose, pH 7.3 with KOH. Borosilicate glass pipettes (1.5 OD, 0.85 ID, Sutter Instrument Company, Novato, CA) were filled with pipette buffer solution (in mM: 130 NaCl, 5 KCl, 10 HEPES, 10 TEACl, 1 CaCl_2_, 1 MgCl_2_, pH 7.3 with NaOH. In a small dataset obtained with pipettes with a resistance of less than 3 MΩ peak current was strongly correlated with pipette size and few patches had zero channels (**Figure S2D**); we therefore restricted our final dataset to pipettes between 3-6.5 MΩ Negative pressure was applied through the patch pipette with an amplifier-controlled high-speed pressure clamp system (HSPC-1; ALA Scientific Instruments, Farmingdale, NY). All pressure protocols were preceded by a +5 mmHg prepulse (+60 mV recordings, 2 seconds; −80 mV recordings, 5 seconds) to remove resting tension due to the gigaseal (Lewis and Grandl 2015).

### Quantification and Statistical Analysis

All data were analyzed and final plots were generated using Igor Pro 8.02 (Wavemetrics). For currents measured during ramp protocols, current was binned for each pressure (1 mmHg increments) and any linear changes in leak prior to the first channel opening were manually identified and subtracted off prior to further analysis. Current-pressure curves were fit with a Boltzmann function

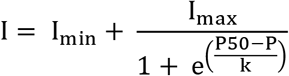

where I_max_ is the maximal current, P is pressure, P_50_ is pressure of half-maximal activation, and k is the slope factor. The number of channels in each patch was calculated by dividing I_max_ by the mean single channel current from all patches with precisely one channel (i.e., Imax during the ramp phase for patches with only one visually identified discrete opening, **Figure 1**) or from single channel openings during the prepulse (step recordings with overexpressed channels, **Figure 4**). To generate idealized currents (**Figure 2**), the number of channels (n) in each patch, as well as the time of openings, were manually identified. Patches with larger numbers of channels (3-5) are underrepresented due to the increased difficulty in identifying discrete openings. For currents measured during step protocols, to reduce artifacts from noise, the peak was measured from a 0.5 ms rolling average around the true peak and the steady-state current was calculated as the mean current during the last 10 ms of the step.

Single-channel currents during the +5 mmHg prepulse were filtered offline (1 kHz) using Fitmaster v2×90.2 and baseline-subtracted using QuB online software (Nicolai and Sachs 2013). Single channel amplitudes during the prepulse were measured by generating all-points histograms with binning calculated using the Freedman-Diaconis method and an optimal bin width of 2*IQR(x)/N^1/3^, where IQR is the interquartile distance, N is the number of observations, and the bins are evenly distributed between the minimum and maximum values. Binned data were then fit with a double-Gaussian equation of the form

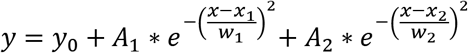

where y_0_ is the baseline current, A_1_ and A_2_ are the peak amplitudes, x_1_ and x_2_ are the centers of the fits, and w_1_ and w_2_ their respective widths. The difference between x_1_ and x_2_ reflects the difference between mean current in the open and closed state and was used to calculate single channel current i.

Open probability during the prepulse was calculated as the mean current (i*n*P_o_) divided by the maximum current elicited for that patch at saturating pressures, assumed to be P_o_ = 1 (i*n). Data are reported as mean ± SD unless otherwise indicated. For box and whisker plots, boxes represent median and first and third quartiles.

### Thomas Clustering Model

A model for spatial distribution of Piezo1 ion channels was built using a Thomas point process in IGOR Pro 8.02 (Wavemetrics) (Baddeley 2015). Parent discs were randomly distributed in a 50 μm x 50 μm arena. Each parent disc was then randomly assigned a varying number of daughter points. For a truly random distribution (mean cluster = 1) every parent disc was assigned precisely 1 daughter point. To introduce a slight propensity for channels to cluster, the number of daughter points were drawn from a Poisson distribution centered at 1, such that the mean cluster size (for populated clusters) was 1.6. To introduce a stronger propensity to cluster, the number of daughter points were drawn from a Poisson distribution centered around 5 (mean cluster = 5); **Figure 3A**. The densities of parent discs and daughter points were adjusted such that their product, and thus the final number of daughter points in the arena, was equivalent to that of the wild-type Neuro2A dataset (1.75 channels/μm^2^). Daughter points were then distributed in each parental disc according to a Gaussian distribution with an average distance of sigma = 50 nm from the center of the disc, or roughly the distance of three Piezo footprint decay lengths. The arena was then sampled with 1,000 circles whose centers were uniformly distributed within the arena. To replicate variability in pipette size/patch dome size as source of noise, the mean radius of all circles was 0.8 μm (see below) with a standard deviation of 0.15 μm, which corresponds to our mean pipette size of 4.35 MΩ and standard deviation of 0.8 MΩ (**Figure S3**). Finally, the number of daughter points (“channel counts”) within each circle was then counted. To further account for variability in transmembrane potentials as a source of noise, we translated channel counts into current amplitudes using our experimentally obtained normal distribution of single channel currents, which yielded a mean single channel current of 0.98 pA and standard deviation of 0.22 pA (**Figure S2F**). Current amplitudes were then reverted into channel counts and used to generate a channel count histogram. Nearest neighbor distributions were obtained from the same simulated distributions as above, by calculating a distance matrix between all channel centers and taking the minimal distance for each row.

### Calculation of channel footprint and density

The characteristic decay length (λ) of membrane deformation induced by the large curved dome of Piezo1 is estimated to be 14 nm (Haselwandter and Mackinnon, 2018). We put an upper bounds of possible Piezo energetic interactions due to membrane curvature beyond the dome itself as 3x this decay length; when adding this to the radius of the channel dome, this yields an approximate total membrane footprint of (10 nm + 3*14 nm ~ 50 nm).

From our previous data ((Lewis and Grandl 2015); **Figure S3**) we estimated that patch domes from pipettes 3-6 MΩ have an approximate surface area of 2 μm^2^. If the patch dome is approximated as a flat circle, this corresponds to a pipette radius of 0.8 μm, which was used for simulation of Piezo1 spatial distribution in the Thomas point process model. For native expression in Neuro2A cells, our average channel count was 3.5 channels, which corresponds to an average density of 1.75 channels/μm^2^. For overexpression of Piezo1 in Neuro2A-p1ko cells, our average channel count was 35 channels, which corresponds to an average density of 17 channels/μm^2^. For the maximal current amplitude we observed (160 pA = ~80 channels), with an estimated in-plane area of 400 nm^2^ (Wang et al. 2019), an estimated footprint of 7,800 nm^2^ (calculated from a footprint of 50 nm (Haselwandter and MacKinnon 2018)) and a patch dome surface area of 2 μm^2^ approximately 2% of the membrane surface would be covered by Piezo channels and 31% of the surface covered by Piezos+corresponding footprints.

---

**Supplemental Figure 1.**
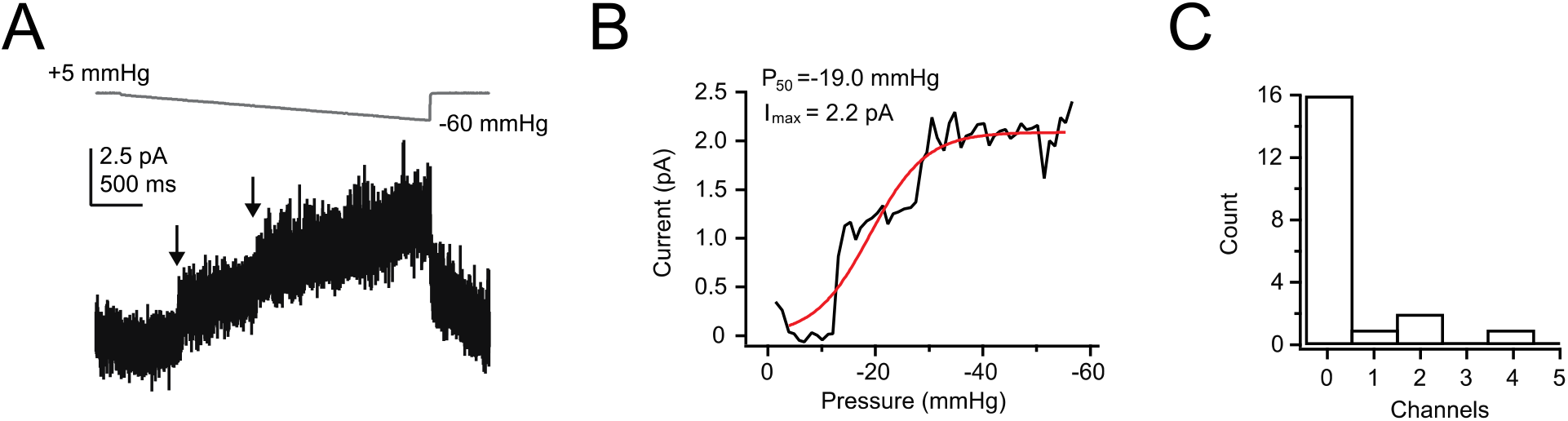
HUVECs express low levels of Piezo1. **A**, Pressure protocol (gray) and current (black) from a cell-attached patch from a wild-type HUVEC cell. Pressure was increased stepwise from +5 mmHg to −60 mmHg in 1 mmHg steps, 50 ms per step. Pressure was clamped at +5 mmHg for 2 seconds prior to the step protocol to remove resting tension from the patch. Holding potential was +60 mV to minimize inactivation. **B**, Current-pressure relationship for the same patch. Mean current was calculated for each step, in 1 mmHg increments, and plotted as a function of pressure. **C**, Histogram of number of channels per patch. n = 20 patches.

**Supplemental Figure 2.**
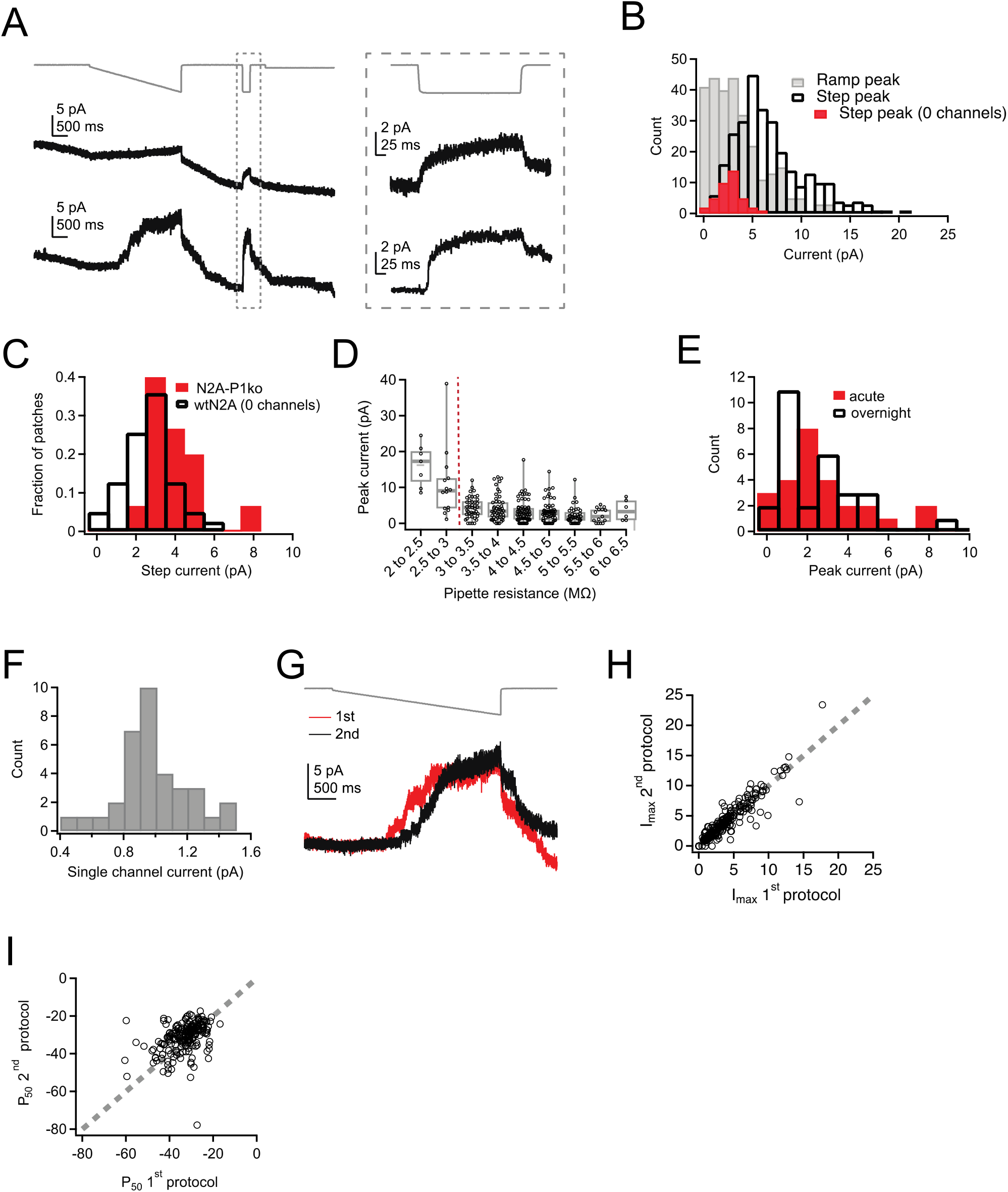
Ramp-like protocol effectively measures I_max_ in Neuro2A cells. **A**, Pressure protocol (gray) and current (black) from two cell-attached patches from wild-type Neuro2A cells, one with zero channels (top) and one with many channels (bottom). Note the step protocol (inset on the right) elicits a small current in the patch with zero channels, despite no evidence of discrete channel openings during the ramp protocol on the left. **B**, Histogram of peak currents elicited by the ramp (gray) and step (black) portions of the protocol. Ramp peak currents are leak subtracted according to the protocol in Figure 1; step peak currents are therefore right shifted due to the inability to subtract the leak current. n = 281 patches. Red, step peak currents for patches with zero channel openings identified during the ramp portion of the protocol (n = 33 patches). **C**, Normalized histograms for peak current elicited by the step protocol for patches from wild-type Neuro2A cells with zero channels (n = 33 patches) and patches from Neuro2A-p1ko cells (n = 15 patches). **D**, Peak current during the ramp protocol as a function of pipette resistance. Only patches made with pipettes with a resistance of > 3 MΩ red dashed line) were used in the final analysis. **E**, Histogram of peak current amplitudes for acutely plated or day-matched overnight plated wild-type Neuro2A cells (see Methods). Acute, n = 26 patches; overnight, n = 28 patches. **F**, Histogram of single-channel current amplitudes, measured from the I_max_ values of patches that had precisely one channel. n = 35 patches. **G**, Pressure protocol (gray) and current for two protocols run successively on the same patch. **H**, I_max_ for the second protocol as a function of I_max_ for the first protocol from the same patch. Gray dashed line represents unity. **I**, P_50_ for the second protocol as a function of P_50_ for the first protocol from the same patch. Gray dashed line represents unity.

**Supplemental Figure 3.**
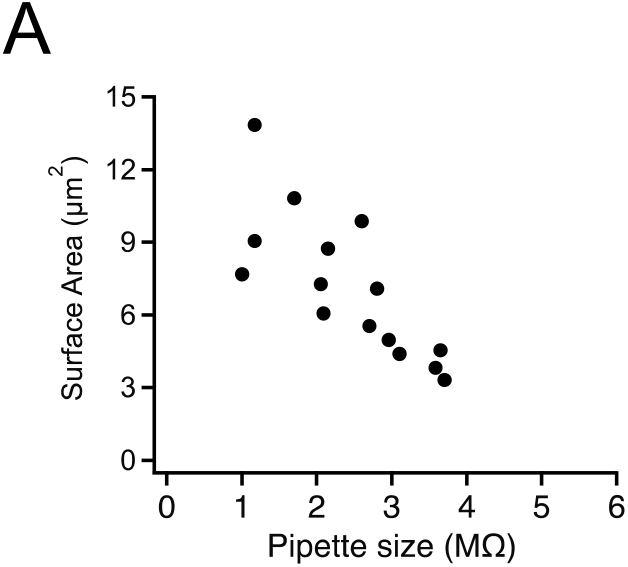
Estimation of patch dome size as a function of pipette resistance. **A**, Surface area as a function of pipette resistance for cell-attached patches from HEK293t cells expressing mouse Piezo1. Data re-analyzed from (Lewis and Grandl 2015).

**Supplemental Figure 4.**
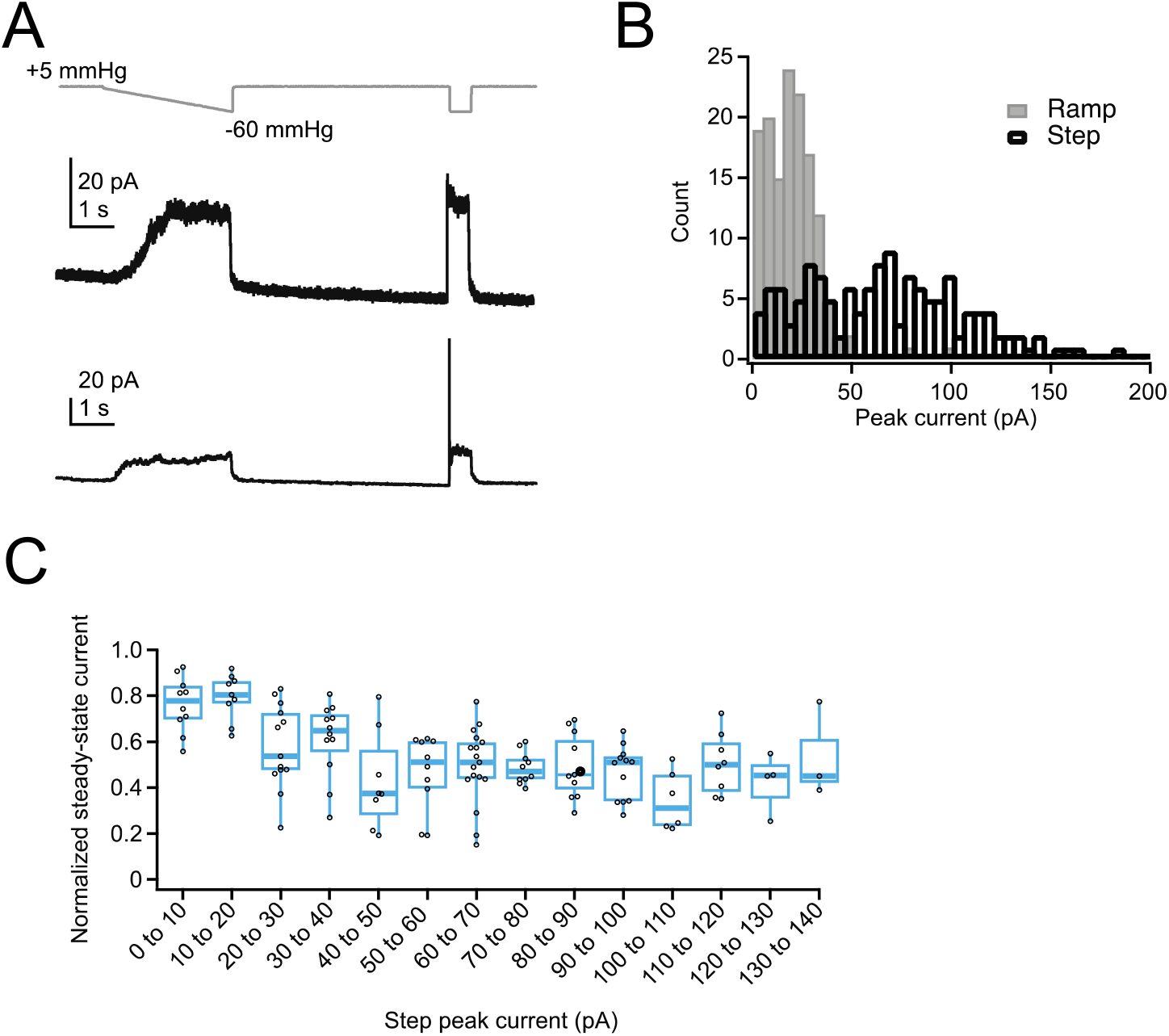
Ramp-like protocol induces substantial inactivation in overexpressed Piezo1 channels. **A**, Pressure protocol (gray) and current (black) from cell-attached patches from two Neuro2A-p1ko cells overexpressing mouse Piezo1. Top patch has minimal inactivation during the step portion of the protocol, bottom patch has substantial inactivation during the step portion of the protocol. **B**, Histogram of peak currents elicited by the ramp (gray) and step (black) portions of the protocol. Note that the ramp protocol substantially underestimates the total peak current due to inactivation. n = 138 patches. **C**, Normalized steady-state current, measured during the last 10 ms of the step pulse, as a function of peak step current.

**Supplemental Figure 5.**
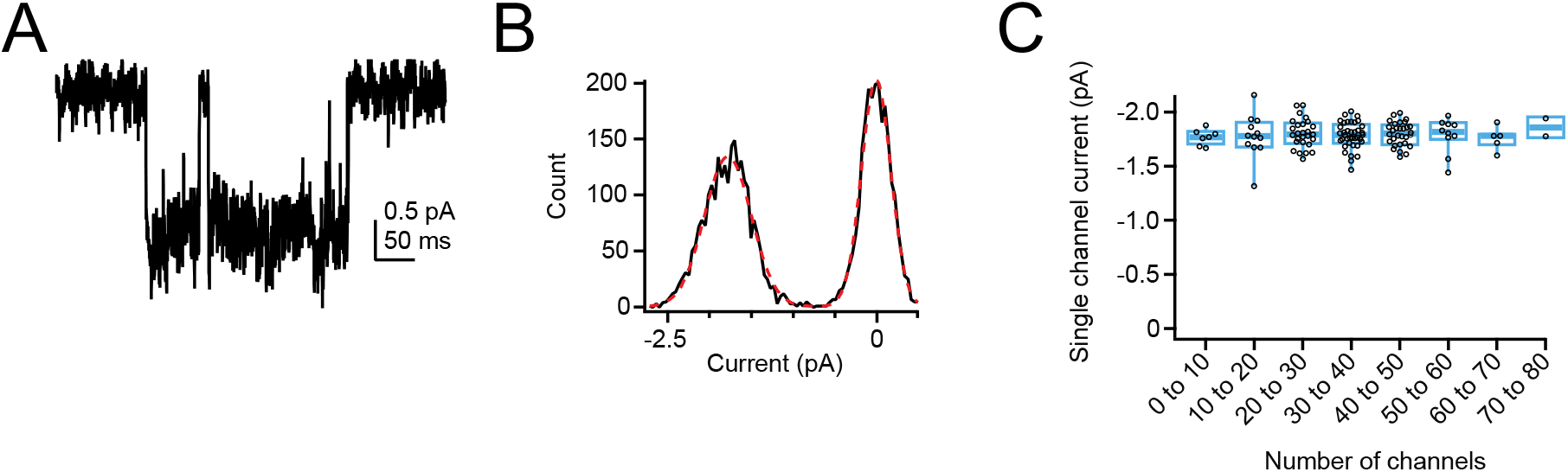
Single-channel current measurements from overexpressed Piezo1. **A**, Single-channel current from a cell-attached patch from a Neuro2A-p1ko cell overexpressing mouse Piezo1. Pressure was +5 mmHg and voltage was −80 mV. **B**, All-points histogram generated from current sweep in A. Red line is a double-Gaussian fit to binned data. **C**, Single-channel current as a function of number of channels in the patch. n = 133 patches.

**Supplemental Figure 6.**
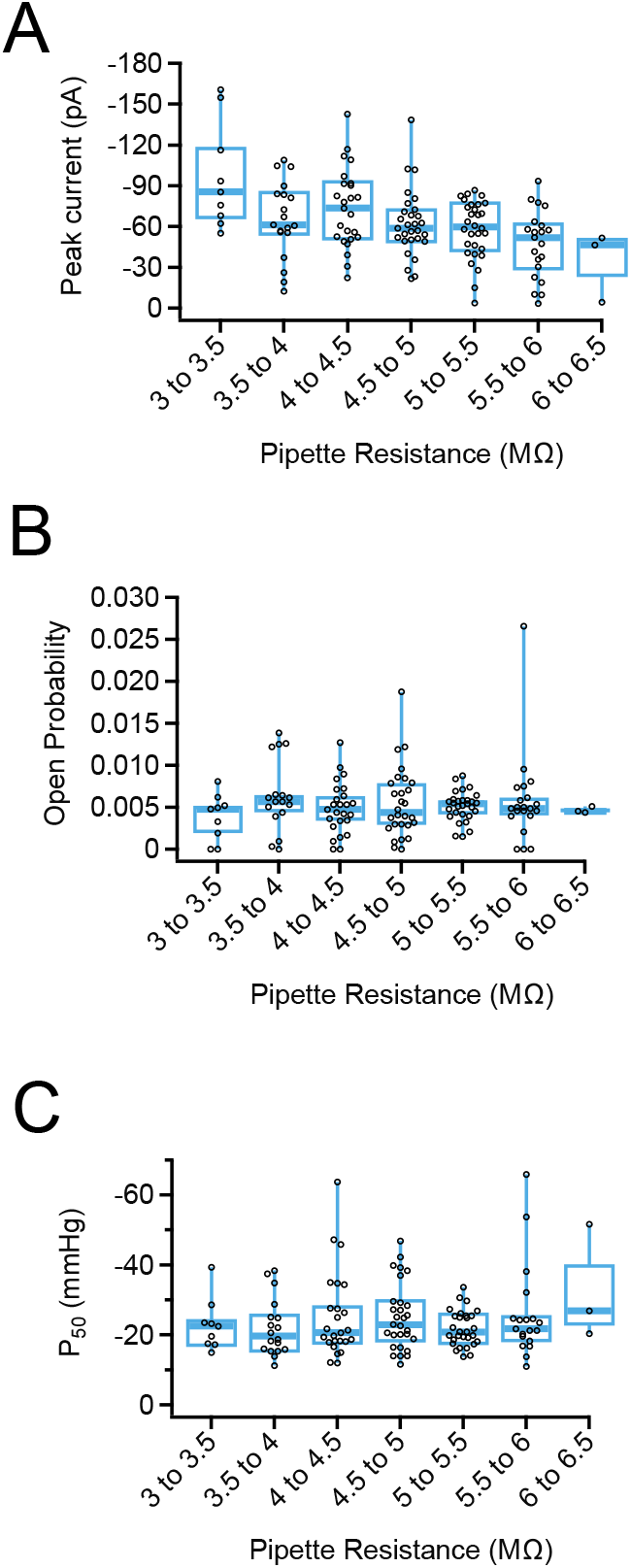
Pipette resistance does not affect Piezo1 open probability or pressure sensitivity. **A**, Peak current as a function of pipette resistance for Neuro2A-p1ko cells overexpressing mouse Piezo1. **B**, Open probability, measured during the +5 mmHg prepulse from the protocol in Figure 4B, as a function of pipette resistance. **C**, P_50_ measured from sigmoidal fits to current-pressure relationships from the protocol in Figure 4B as a function of pipette resistance.

**Supplemental Figure 7.**
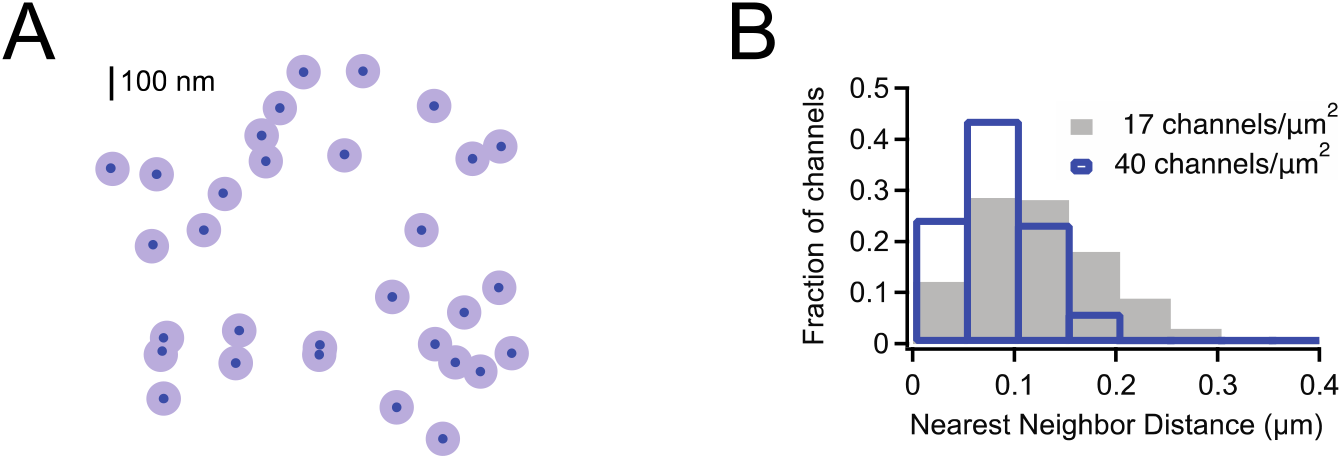
Estimation of nearest-neighbor distances for overexpressed Mouse Piezo1. **A**, Representative channel distribution generated using a Thomas point process with overall densities equivalent to that of the highest density we see with overexpression of mouse Piezo1 in Neuro2A-p1ko cells (~40 channels/μm^2^). Dark blue circles are the projected area of the Piezo footprint to scale; light blue circles are ~3x the membrane footprint for each channel (radius = 50 nm). **B**, Histogram of nearest-neighbor distances for densities representing the mean density in our overexpression dataset (~17.5 channels/μm^2^) and the maximum density (~40 channels/μm^2^).

